# A Space-Time Hidden Markov Model for In Vivo NanoScale Synaptic Plasticity Tracking

**DOI:** 10.1101/2025.10.10.681691

**Authors:** Shashwat Kumar, Gabrielle I. Coste, Dasun Premathilaka, Richard L. Huganir, Austin R. Graves, Adam S. Charles, Michael I. Miller

**Affiliations:** Department of Biomedical Engineering, Johns Hopkins University, Baltimore, MD, USA; Department of Neuroscience, Johns Hopkins University, Baltimore, MD, USA; Kavli Neuroscience Discovery Institute, Johns Hopkins University, Baltimore, MD, USA

## Abstract

Synapses are the fundamental unit of neural connectivity, exhibiting dynamic functional and structural changes that enable the brain to learn, adapt, and form memories. Recent advances in fluorescent labeling of endogenous proteins offer an opportunity to image synaptic strength in vivo and study mechanisms underlying adaptive neural computation. Studying synaptic dynamics requires tracking signals of small, densely packed synapses over days as they change in size, position, and intensity between imaging sessions, and may even appear or disappear. Associating *>*50,000 dynamic, submicrometer particles across time is difficult, even for state-of-the-art algorithms. Moreover, most algorithms assign equal weight to the lateral (XY) and noisier axial (Z) dimensions, reducing performance. To address these challenges and accurately track synapses in vivo, we developed SynTrack. We formulate tracking as a Maximum *A Posteriori* estimation problem that identifies the K most likely disjoint paths in a Hidden Markov Model, solved using min-cost circulation optimization. An anisotropic uncertainty model accounts for poorer axial resolution, and a fully temporally connected spatio-temporal graph overcomes long-term occlusions. SynTrack achieves a mean displacement of 0.50 ***µ***m with a Multiple Object Tracking Accuracy (MOTA) score of 89.8%, on par with expert annotators but with substantially increased speed and scalability. In a large-scale volume imaged over two weeks, SynTrack reconstructed 74,000 synapse trajectories detected in 4.9 out of 8 imaging sessions on average, with 18,000 synapses tracked in at least seven sessions. We present a state-of-the-art algorithm capable of high-fidelity longitudinal tracking of individual synapses in behaving mice at an unprecedented scale.

## 1 Introduction

Synapses are densely packed sub-micron structures that connect pairs of neurons and enable propagation of electrical signals across vast ensembles of neurons via synaptic neurotransmission. As animals form memories, adapt to their environment, and learn to perform new tasks, synaptic connections strengthen or weaken (by adding or removing neurotransmitter receptors, respectively), and new synapses can be created or deleted. This process of dynamic remodeling of synapses is called synaptic plasticity, which has long been a central focus in neuroscience and is thought to underlie many forms of learning, memory, and behavioral adaptation. AMPA-type glutamate Receptors (AMPARs) are known to be centrally involved in synaptic plasticity, specifically in long-term potentiation and depression [1, 2]. Given the critical role plasticity plays in the brain, the ability to track its timecourse and spatial patterning in living animals would significantly advance our understanding of this central function of the brain, allow us to observe how synaptic deficits are manifested in neurological disease, and pave the way to understand new learning rules in artificial systems. However, studies of the computational impacts of plasticity have largely been theoretical [3–6] due to the difficulty of observing the dynamics of hundreds of thousands of synapses in vivo across long time-scales. Only recently have microscopy, biological, and algorithmic tools advanced to the point of resolving submicrometer-sized structures across days to weeks, thus opening the door to such studies.

Nevertheless, the difficulties posed by in vivo imaging of micron-sized synapses in living animals present significant resolution and motion challenges. Previous work imaging changes in AMPAR expression during learning tasks have been restricted to observing a sparse set of synapses (fewer than *≈*1000 synapses per mouse [7]) or ex vivo studies using confocal microscopy in postmortem tissue [8]. More recently, new transgenic mouse lines [9], combined with multiphoton imaging [10] and novel algorithms for image enhancement [11] and segmentation [12], have reported longitudinal imaging of hundreds of thousands of synapses in vivo (Fig. 1). Such datasets finally provide faithful observations of large-scale synaptic changes, though identifying individual synapses and tracking their changes in strength over time (i.e. synaptic plasticity) with high fidelity remains a significant hurdle.

**Figure 1:**
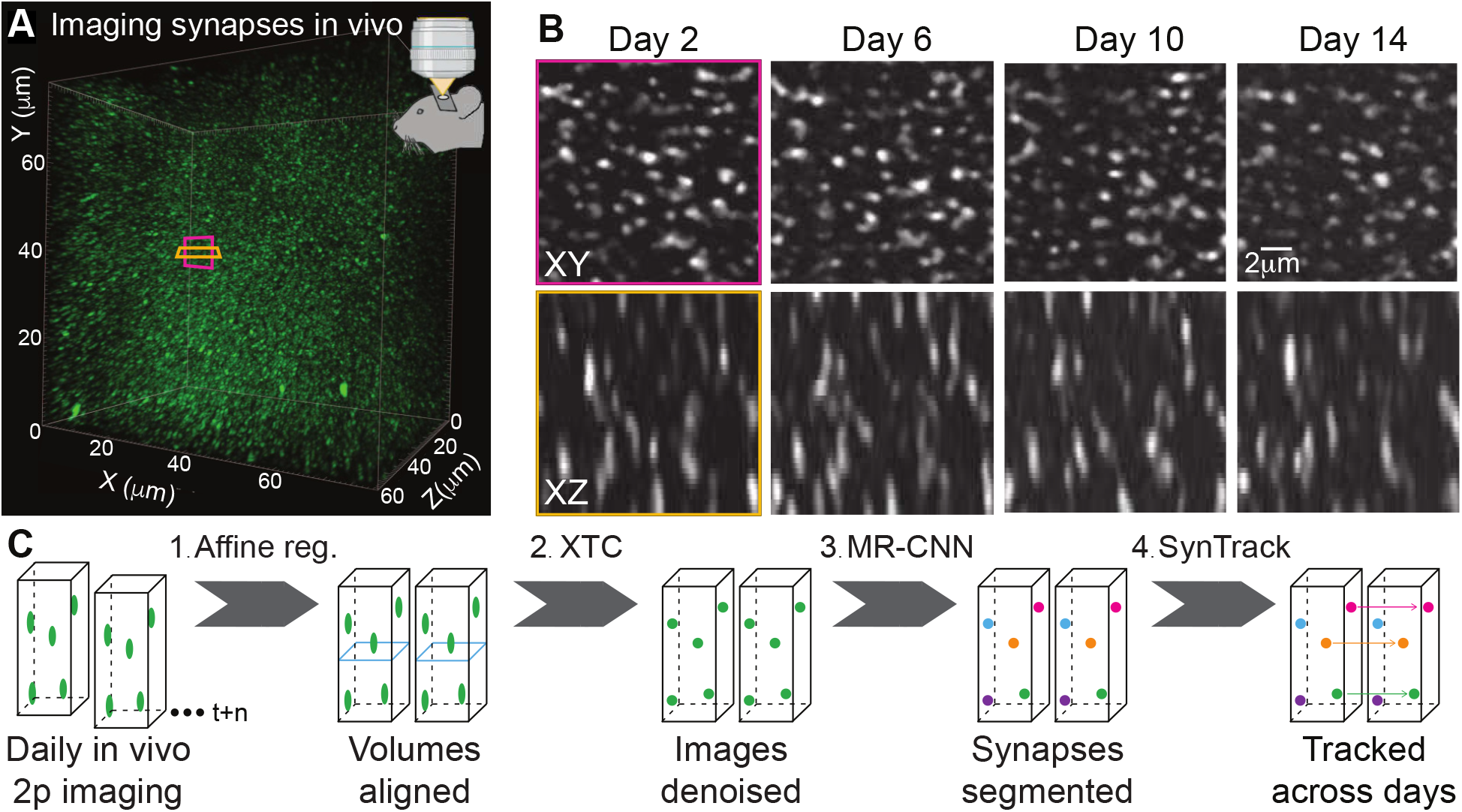
The synapse tracking problem. (A) In vivo imaging volume comprising 50,000 cortical synapses of a SEP-GluA2 mouse. Each green blob is a single excitatory synapse. Pink and orange boxes respectively represent the location of XY and XZ planes shown in (B). (B) Longitudinal images after affine registration. Sum projection of 4 slices (1.33 *µ*m total thickness). SynTrack must take into account larger synapse location uncertainty in Z caused by the microscope PSF. (C) Synapse imaging, detection, and tracking pipeline: after longitudinal 2-photon (2p) imaging in vivo, imaging volumes are aligned with affine registration and denoised with XTC [11]. Synapses are then detected with MR-CNN [12]. SynTrack links synapse detections between days to create longitudinal tracks.

Several key challenges make tracking densely labeled synapses in vivo difficult. First, the geometry of in vivo imaging and limitations on the objective lens make the Point-Spread Function (PSF) have coarser resolution along the axial dimension (Z) compared to XY planes. The asymmetric resolution produces an anisotropic image, which results in higher axial uncertainty of synapse structure and position that must be taken into account in a tracking algorithm. Second, while modern transgenic mouse lines that label *all* excitatory synapses provide a unique opportunity to understand how synaptic plasticity drives cognition, the density of labeled synapses presents a challenge for matching and tracking hundreds of thousands of individual synapses across imaging sessions (Fig. 1a). Third, individual synapses lack distinctive features, and thus identity information is principally derived from their spatiotemporal location relative to other synapses (Fig. 1b) rather than from distinguishing shape or textures. Finally, the high dimensionality and limited individuality of synapse shape is further exacerbated by the limited Signal-to-Noise Ratio (SNR) inherent to imaging small objects in scattering tissue. The limited SNR can impact the accuracy of localizing or even detecting a given synapse on one or more sessions. Thus, a tracking algorithm must be able to re-identify missing synapses between days. Furthermore, as data are acquired by imaging large volumes of brain tissue, synapse tracking must be performed in 3D.

Due to the above challenges, published synapse tracking has largely focused on manually tracking relatively small numbers of synapses from sparsely labeled samples by identifying their location relative to an anatomical anchor, e.g., a fluorescently labeled dendritic spine [7]. Brain-wide imaging and tracking of extremely large numbers of neurons, however, has been achieved [13, 14], lending support to the feasibility of tracking similar numbers of synapses in living mice. There is a rich body of literature in the field of in vivo biological cell tracking across time series, including simple nearest neighbor methods [15], greedy pairwise matching between frames [16, 17], and more recent approaches employing deep-learning algorithms for linking [18]. In some cases, deep-learning approaches have achieved state of the art segmentation but often demonstrate poor generalizability to datasets that are “out of distribution” with respect to training data. A recent 10-year benchmark on the cell tracking challenge [19] reported that traditional approaches for data association can be as effective as deep learning.

To enable high-throughput synaptic tracking, we introduce SynTrack, a probabilistic framework for reconstructing synapse trajectories from longitudinal imaging data. At the core of SynTrack is a novel formulation of synapse tracking as inference in a directed acyclic graph (DAG) structured hidden Markov model (HMM) over space-time, where latent states represent synapse locations and detections are treated as noisy observations. This formulation explicitly captures anisotropic spatial uncertainty arising from the optical point spread function, as well as random detection censoring due to missed or spurious detections. By allowing transitions across arbitrary temporal gaps, the model naturally handles missing detections and enables robust re-identification of synapses across imaging sessions.

We estimate synapse tracks by computing the maximum a posteriori (MAP) solution under this HMM via minimum cost circulation optimization. This yields a scalable algorithm capable of aligning hundreds of thousands of synapses, while remaining robust to dense clutter, low signal-to-noise ratio, and partial observations.

## 2 The SynTrack Method

We represent each detected synapse by its spatial location and imaging session. We then model detection dropout, false positives, and anisotropic spatial uncertainty within a hidden Markov framework over candidate synapse trajectories. Specifically, our statistical model is a HMM in which the trajectories are paths through ℝ^3^ *× T* . To estimate trajectories under this model, we pose synapse tracking as a minimum cost circulation problem in the space of detected synapses. To ensure proper tracking, we preprocess the sequence of image volumes by aligning each session to a common template coordinate system using affine registration. Specifically, each collected volume is mapped to the template with an affine transformation that corrects for large-scale motion common to in vivo imaging, including microscope misalignment, breathing, and blood-pressure-related shifts. The resulting registration transforms are then applied to the detected synapse coordinates before tracking.

### 2.1 Hidden Markov Probabilistic Directed Acyclic Graph Models

Let *𝒟*= {*d*_*i*_} _*i∈I*_ denote the set of detections, indexed by *I* with cardinality |*I* |= *N* . Each detection is associated with a spatial location *x*_*i*_ *∈* ℝ^3^ and timestamp *t*_*i*_ *∈* {1, …, *T*} . We consider two observation models:

In the simplest setting, the detections represent spatiotemporal coordinates together with the CNN-estimated confidence *θ*_*i*_ *∈* [0, 1] that the detection corresponds to a true synapse:

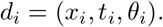

Alternatively, detections can represent a set of pixel intensity values observed inside each segmentation volume

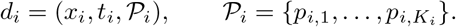

The detections in *𝒟* arise from the pipeline in Figure 1c, which uses CNN-based machine learning to denoise [11] each imaged volume and then subsequently identifies hundreds of thousands of putative synapse segmentations [12] in the sequence of observed imaging volumes.

We define the state as the spatiotemporal location of a synapse track, i.e., *s*_*i*_ = (*x*_*i*_, *t*_*i*_). Each detection *d*_*i*_ is associated with a state *s*_*i*_. The state space of synapses is constructed as space time locations augmented with a start and terminal state:

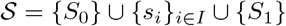

We model transitions between states in *𝒮* using a stochastic transition kernel

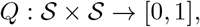

where

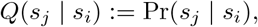

with

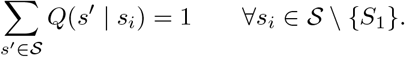

We define artificial timestamps for the source and terminal states:

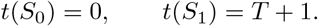

To enforce temporal ordering, transitions are only permitted strictly forward in time:

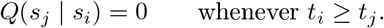

The time ordering places our probabilistic models into the class of directed HMMs [20], i.e., the ordering of the arrows (in contrast to general random fields) imply that descendents are conditionally independent given their parents (see Fig. 4).

### 2.2 Bayesian Model and MAP Estimation

#### 2.2.1 Synapse Track parameterization

We formalize the space of probabilistic models for synapse tracks as directed acyclic graphs. A synaptic track is a cluster forming a linear graphical connection of the “hidden” time-ordered sequence of spacetime locations in ℝ^3^ of synapses of identical identity. These clusters are always a partial covering of the full state space *𝒮*. The tracks are generated from collections of sites in our sampling lattice of synapse locations associated to identified putative synapses *𝒟*.

We identify tracks with the time-ordered index sequences:

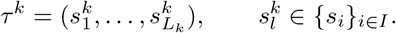

such that

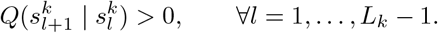

Thus, each track corresponds to a path in the directed acyclic graph induced by the transition kernel *Q* and is inherently time-ordered. The start and end states are fixed and do not contribute to the degrees of freedom.

A configuration of *K ≥* 1 tracks with 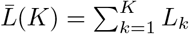 is written as:

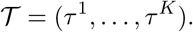

Our space of tracks forms a partial or complete covering of the indexed set of hypothesized fluorescence peaks *𝒮*, the size of which |*𝒮* |= *N* + 2 determines the complexity of our problem. We assume a partial covering by tracks since some detections are false alarms that do not necessarily correspond to synapses.

#### 2.2.2 The Space of Observed, Detected Synapses

Two sources of uncertainty are modeled impacting how the tracks are inferred from the data: (i) the synapse detections in a session are modeled as noisy observables forming our likelihood *𝓁* on *d ∈ 𝒟*scoring our confidence of the detections, and (ii) the spatial uncertainty in synaptic location between days of imaging, modeled with our prior distribution *p*_Σ_ on *𝒯*. The parameter Σ represents the spatial variation of the latent tracks.

These together define our cost for Bayesian maximum a-posteriori estimator (MAP) which we generate via min-cost circulation optimization. Viewing our track parameterizations as random defines our posterior probability laws defining the MAP:

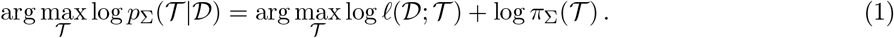

#### 2.2.3 Likelihood Model

Our likelihood models the observed detections *d*_*i*_ *∈ 𝒟* as a probabilistic function of *𝒯*. The subset of the detections covered by the tracks is defined as the *foreground*, with the complement defined as the *background*, i.e., the false detections.

The likelihood of the foreground detections *d*_*i*_ *∈ 𝒟* is modeled as independently distributed given *𝒯* with a spatial field of parameters *{*0 *≤θ*_*i*_ *≤* 1, *i ∈ I }* representing the confidence of the foreground.

We define the set of indices corresponding to detections covered by the tracks as:

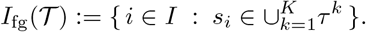

This allows us to define the detection likelihood in terms of the foreground background likelihood:

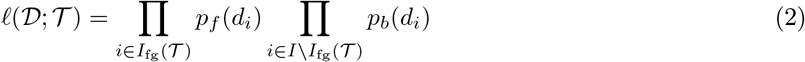

We use the Bernoulli likelihood *p*_*f*_ (*d*_*i*_) = *θ*_*i*_, *p*_*b*_(*d*_*i*_) = 1 *−θ*_*i*_. Alternatively, a histogram estimate for foreground and background voxel intensity distributions can also be used. The key idea of our likelihood model is that the tracks should “cover” higher confidence object detections while “pruning” lower confidence ones.

#### 2.2.4 The prior model on track continuation

The transition kernel defines the prior probability of a single track *τ* = (*s*_1_, …, *s*_*L*_) of length *L*:

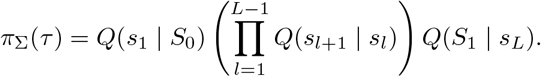

Here, *Q*(*s*_1_ |*S*_0_) denotes the probability of initiating a track at state *s*_1_, while *Q*(*S*_1_ |*s*_*L*_) denotes the probability of terminating the track at state *s*_*L*_.

Ignoring the mutual exclusion constraints, the prior score over a configuration of K tracks factorizes as

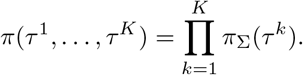

### 2.3 The MAP Estimator in its Constraint Space

Our MAP estimate 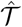 of the track clusters maximizes the posterior of Equation (1) subject to the constraints. Our prior model has only first order Markov structure as written, but the constraints increase the graphical conditional probability dependence. We condition on the number of *K ≥* 1 tracks of lengths *L*_*k*_, *k* = 1, …, *K*. We assume each parse sequence is time-ordered without cycles so that no synaptic time-position within any track is revisited. Multiple tracks are assumed to be non-overlapping disallowing two tracks from covering the same detection.

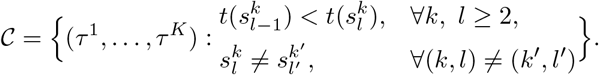

#### Definition 1

(MAP ESTIMATOR).

*Let 𝒯* = (*τ* ^1^, …, *τ* ^*K*^) *represent a track configuration, the MAP satisfies*

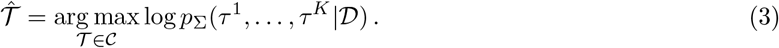

For *K*-track configurations *𝒯*= (*τ* ^1^, …, *τ* ^*K*^) coupling between tracks is implemented through the min-cost circulation graph.

### 2.4 Minimum Cost Flow Circulation Graph Reparameterization

The optimization for maximizing our cost function is an integer linear programming problem. We solve it using min-cost circulation algorithms by re-stating it as an optimization over a graphical representation *G* = (*V, E*) of the HMM-DAG: Given the set of indices *I*, we duplicate them and call the set *I*^*\**^. The set of vertices is:

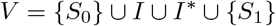

Define the set of edges:

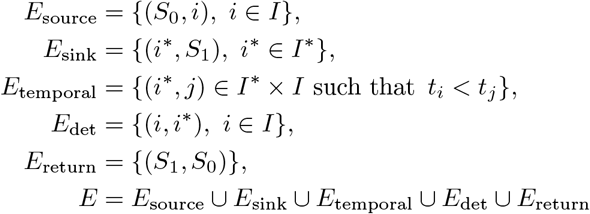

To enforce the constraints on the graph of a single connected cluster we define the set of incoming and outgoing edges for each graph node:

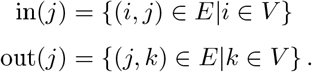

We associate two variables with each edge. The first is a nonnegative integer-valued flow variable *f* : *E→* ℤ_*≥*0_, where *f* (*e*) denotes the amount of flow assigned to edge *e ∈ E*. The second is a cost of using that edge *c* : *E→* ℝ which translates the probability of a transition into a non-negative cost of the associated edge in the min-cost optimization.

We define an edge capacity variable

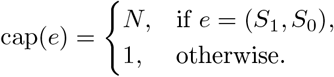

The objective in Equation (3) is formulated as a min-cost circulation problem.

The optimization formulated this way is an integer linear programming problem, rather than a dynamic programming generalization of the Viterbi algorithm as proposed in [21]. Choosing the costs as specified in Statement (1) below gives us our MAP estimate.

#### Lemma 1

(Path Disjointness of Flows). *All feasible flows f on G obtained as solutions to Algorithm 1 decompose into vertex-disjoint paths from S*_0_ *to S*_1_.

*Proof*. Each detection node *i ∈ I* is split into *i → i*^*\**^ with unit capacity 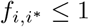. By flow conservation,

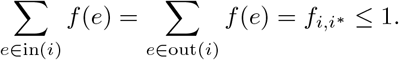

#### Algorithm 1

Min-Cost Circulation Optimization for the Multi-Track Problem

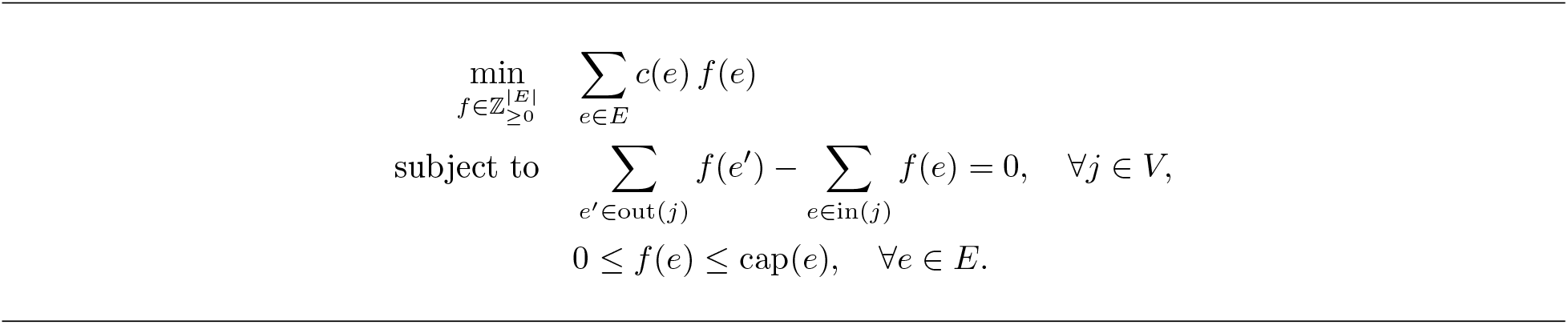

Thus, at most one unit of flow passes through each node, preventing merging or splitting of paths. Since the graph is acyclic (ignoring the return edge), any feasible flow decomposes into directed *S*_0_–*S*_1_ paths, which are therefore vertex-disjoint.

#### Lemma 2.

*Flows f on G are identifiable with tracks* 𝒯 *and vice versa*.

*Proof*. The construction of *G* imposes a time-ordered directed acyclic graph structure. Consider a time-ordered sequence of nodes *τ*^*k*^ = (*s*_1_, …, *s*_*L*_) *∈* 𝒯 . By setting

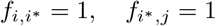

for consecutive states *s*_*i*_ *→s*_*j*_ in the sequence, and including edges from *S*_0_ and to *S*_1_, we can represent each track *τ*^*k*^ as a unit flow along a path in *G*.

Conversely, by Lemma 1, any feasible flow decomposes into vertex-disjoint paths from *S*_0_ to *S*_1_. Each such path defines a time-ordered sequence of states, yielding a track in 𝒯 .

Thus, flows *f* and track configurations 𝒯 are in one-to-one correspondence.

#### Statement 1.

*Set for all e*_*ij*_ *∈ E, i, j ∈ V*,

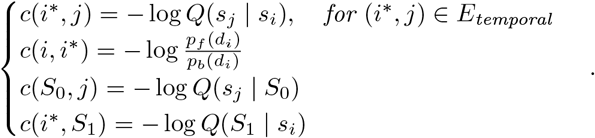

*Then Algorithm 1 gives our Bayes optimum of* (3).

*Proof*. Since each detection index *i ∈I* corresponds uniquely to a state *s*_*i*_ *∈ 𝒮*, we use index notation when defining the flow graph.

Consider the posterior:

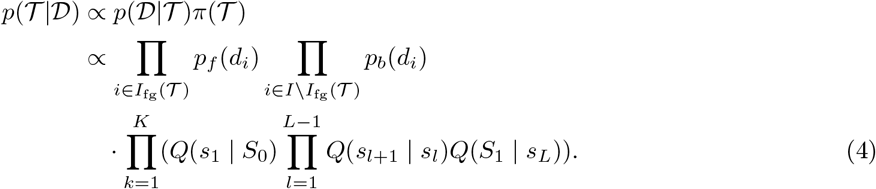

By Lemma (2), we can reparameterize the above objective in terms of the flow variable *f*, yielding

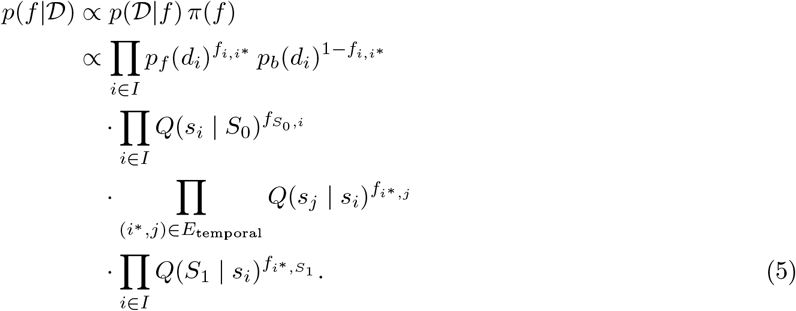

Taking logs of (5) yields the edge weights in (1).

We used the cost scaling algorithm implementation from the Lemon C++ library due to its efficiency. Note that because of total unimodularity, we did not need to formulate this as an integer linear program.

## 3 Results

We tested SynTrack on multi-day two-photon imaging of living mice. The dataset consisted of images of fluorescently labeled synapses from the SEP-GluA2 transgenic mouse line, which enables visualization of all GluA2-containing excitatory synapses across the brain [9, 11]. In this line, endogenous AMPAR GluA2 subunits are tagged with the green fluorophore Super Ecliptic pHluorin (SEP), such that two-photon excitation at 910 nm through a cranial window reveals hundreds of thousands of fluorescent puncta whose intensity (the sum of all voxels within each segmentation) is directly correlated with synaptic strength [9]. Data were collected over a 100 x 100 x 60 *µm* volume for 8 imaging sessions spanning two weeks. All animal procedures were approved by the Johns Hopkins Animal Care and Use Committee and were conducted in accordance with institutional and national guidelines for the care and use of laboratory animals.

We evaluated SynTrack on two complementary in vivo synapse imaging datasets. First, we used a small benchmark volume manually tracked by two expert annotators. This dataset enabled quantitative comparisons using three standard tracking metrics: (1) **#Switches** counting how many times the tracker swaps an object’s identity between timepoints, (2) **IDF1** evaluating how coherently the tracker maintains temporal identity, and (3) Multi Object Tracking Accuracy (**MOTA**), which measures overall tracking accuracy by combining errors like missed detections, false positives, and ID switches [22]. Second, we applied SynTrack to a large-scale volume to assess scalability and demonstrate longitudinal tracking of synaptic dynamics in vivo.

### 3.1 Benchmarking SynTrack against expert annotations

To quantitatively benchmark SynTrack, we first evaluated performance on a small manually annotated in vivo dataset. This benchmark volume contained approximately 100 synapses per imaging session across 8 imaging sessions and was independently tracked by two expert annotators. Because ground-truth correspondences are required to compute standard multi-object tracking metrics, this dataset was used to evaluate identity preservation, tracking accuracy, and agreement with human experts. A critical part of SynTrack is the uncertainty in synapse locations across recording days, which was encoded in the spatial covariance matrix Σ. Accordingly, estimating Σ was an essential step, for which, we used labeled tracks provided by an expert human annotator. We took the difference between consecutive timepoints to estimate the uncertainty in motion between days.

In order to efficiently find pairs of neighbors *x*_*i*_ and *x*_*j*_ that are within *r*^*α*^ Mahalanobis ball of each other, we converted them to euclidean distances by linear transformation 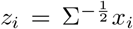 and a CKDTree implementation in scipy.

On this expert-annotated benchmark dataset, tracks generated by SynTrack had a mean spatial displacement of 0.50 *µ*m between consecutive timepoints and were detected on an average of 5.2 out of 8 imaging sessions.

Notably, the temporal and spatial statistics of SynTrack’s output closely matched those derived from expert annotations, suggesting that the method produced biologically meaningful and consistent results, though at vastly improved speed compared to human annotators (a few seconds vs. 6-10 hours for a dataset comprising approximately 100 synapses over 8 imaging sessions).

#### Comparing Trackers

To benchmark SynTrack, we estimated the motion covariance Σ from the ground truth generated by one of our expert annotators. SynTrack achieved a MOTA of 89.8%, an IDF1 score of 83.7%, and only 76 ID switches, closely matching the reference annotator. In comparison, Expert 2 evaluated against Expert 1 achieved a MOTA of 87.8%, IDF1 of 82.7%, and 91 ID switches, indicating greater disagreement. These results show that SynTrack produces trajectories that are on par with two expert human annotators.

We compared SynTrack against standard baselines including Hungarian Matching (adapted from [16]), Nearest Neighbor, and Ilastik tracking with learning. As shown in Table 2, SynTrack outperforms all baselines by a substantial margin, achieving higher MOTA and IDF1 while significantly reducing identity switches. In particular, Hungarian Matching achieves a MOTA of 84.7% and IDF1 of 74.3%, with 114 switches, highlighting the advantage of global data association over greedy approaches. This was observed to be in line with previous findings in cell tracking literature [19] where global data association methods using information from multiple frames tend to outperform greedy methods.

**Table 1:** Average length of tracks, mean distance between consecutive tracked synapses, and average proportion of untracked synapses on each imaging day.

| Tracker | Track length (d) | Pair distance ($\mu\text{m}$) | Untracked (%) |
| --- | --- | --- | --- |
| Expert 1 | 4.9 | 0.532 | 7.5 |
| Expert 2 | 5.3 | 0.564 | 11.6 |
| SynTrack | 5.2 | 0.499 | 4.3 |

**Table 2:** Comparison with classical tracking algorithms and human annotators. Fewer switches is better.

| Method | MOTA (%) ↑ | IDF1 (%) ↑ | Switches ↓ |
| --- | --- | --- | --- |
| Gen-Gauss (geom) HMM | **89.8** | **83.7** | **76** |
| Expert 2 → Expert 1 | 87.8 | 82.7 | 91 |
| Hungarian Matching | 84.7 | 74.3 | 114 |
| Nearest Neighbor | 76.5 | 73.3 | 174 |
| Ilastik Tracking (Learning) | 24.6 | 43.7 | 120 |

One crucial feature of SynTrack was its ability to re-identify “missing” synapses between imaging sessions. In Fig. 3, we compared an XY slice from the ground truth dataset against SynTrack. SynTrack achieved good correspondence with ground truth and was able to reidentify the highlighted synapse that went missing on Day 8 and Day 12.

#### Effect of Tail Modeling and Temporal Priors

Table 3 reveals a clear tradeoff between association precision and temporal consistency within a Gaussian motion model, governed by the tail behavior of the likelihood and the temporal prior. Under the Gaussian assumption, transition probabilities decay quadratically with Mahalanobis distance, allowing long-range associations that can introduce ambiguity in dense regimes. Aggressive truncation (3*σ*) effectively limits this Gaussian tail, yielding the highest IDF1 (84.2%) by suppressing ambiguous long-range associations and improving identity precision. In contrast, moderate truncation (4*σ*–5*σ*) retains more of the Gaussian support, achieving the best MOTA (90.9%) and the lowest number of switches (67), reflecting improved track continuity. Incorporating a geometric temporal prior further regularizes the Gaussian model by penalizing large temporal gaps, consistently reducing identity switches and improving MOTA. Together, these results show that in the Gaussian setting, spatial truncation is necessary for enforcing temporal coherence.

**Table 3:** Effect of penalizing tails with a Generalized Gaussian and a geometric temporal priors on tracking performance (HMM only). Gen-Gauss uses *α* = 2.5, *λ* = 0.5 with a geometric prior *p* = 0.85.

| Model | MOTA (%) | IDF1 (%) | Switches |
| --- | --- | --- | --- |
| Full | 90.7 | 79.4 | 69 |
| Full-Gauss (no geom) | 89.1 | 80.1 | 81 |
| $3\sigma$ | 89.9 | 84.2 | 75 |
| $4\sigma$ | 90.9 | 80.4 | 67 |
| $5\sigma$ | 90.7 | 79.4 | 69 |
| Full Gen-Gauss | 89.8 | **83.7** | 76 |
| Full Gen-Gauss (no geom) | 86.8 | 81.9 | 98 |
| $3\sigma$ | 88.2 | 82.5 | 88 |
| $4\sigma$ | 89.8 | **83.7** | 76 |
| $5\sigma$ | 89.8 | **83.7** | 76 |

The generalized Gaussian model provides a principled alternative to hard truncation, achieving near-optimal IDF1 (83.7%) without requiring explicit threshold tuning. This suggests that generalized Gaussian decay acts as a continuous relaxation of truncation, enabling effective suppression of spurious long-range associations.

### 3.2 Large-scale in vivo synapse tracking

To assess scalability, we next applied SynTrack to a larger in vivo imaging volume containing approximately 50,000 detected synapses per imaging session. Because synapses may be missed, appear, disappear, or only be confidently detected in a subset of sessions, the total number of reconstructed tracks exceeded the number of detections in any single session. SynTrack generated 74,000 total trajectories, with tracks present in an average of 4.9 out of 8 imaging sessions and 18,000 tracks detected in at least seven sessions (Fig. 2). For each synapse within a given imaging session, synaptic strength was defined as the summed fluorescence intensity of all voxels within the automatically segmented puncta. These longitudinal trajectories of fluorescent intensity were used to measure AMPAR content at single-synapse resolution. This enables analysis of synaptic plasticity at a scale not previously achieved in vivo. In Fig. 6, we show dynamics of > 8,000 synapses tracked in all imaging sessions, revealing diverse trajectories of increasing, decreasing, and stable synaptic strength.

**Figure 2:**
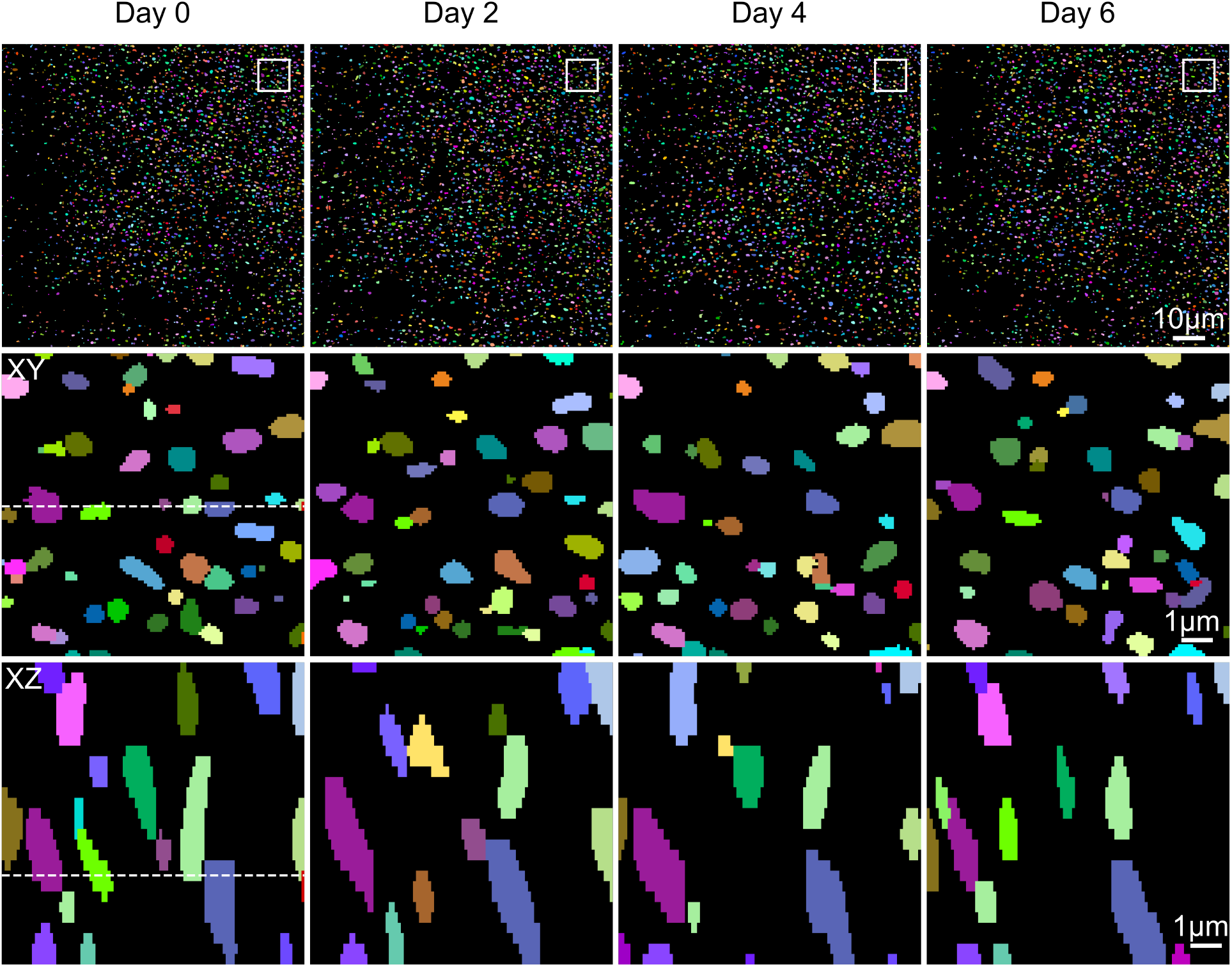
Qualitative Results from SynTrack. Synapse segmentations are color-coded according to their assigned track, with matching colors across columns indicating the same synapse track. White squares in the top row mark the regions shown as zoomed XY images in the middle row. Dotted line highlights the coordinate of the XY and XZ slices.

**Figure 3:**
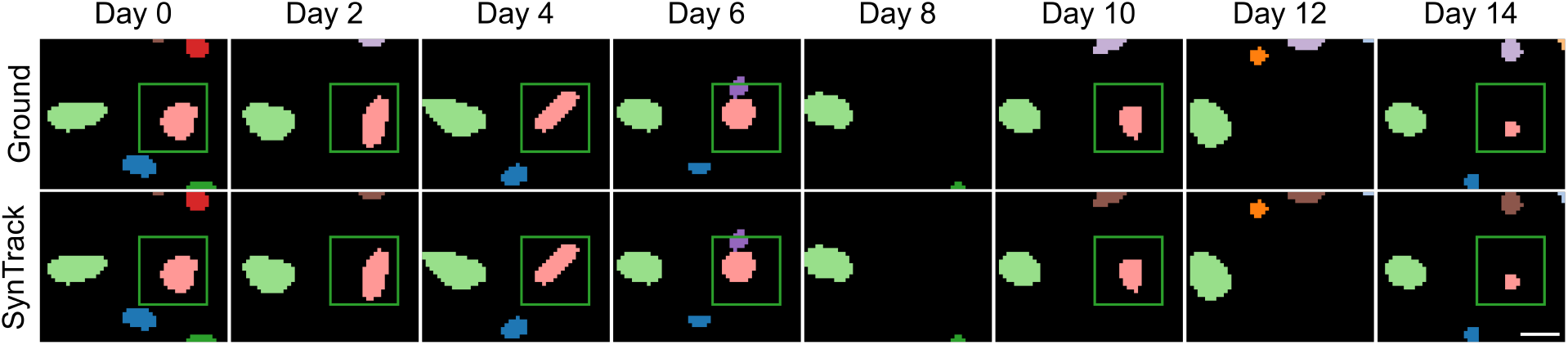
SynTrack successfully re-identifies synapses that go missing and reappear on consecutive sessions (highlighted by green box). Top row shows ground truth tracks labeled by expert annotator. Bottom row shows the corresponding synapses automatically tracked by SynTrack. Synapse segmentations are color-coded according to their assigned track. Scale bar: 1 *µ*m.

**Figure 4:**
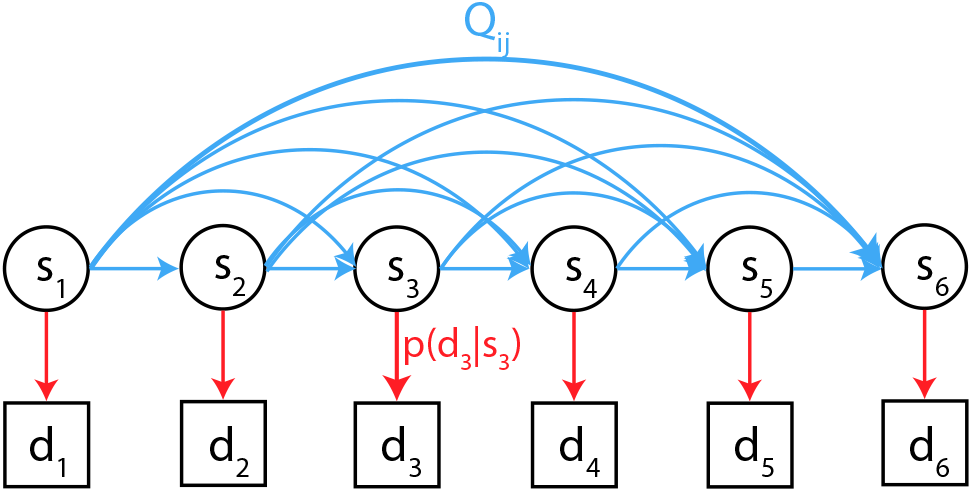
Depiction of our Hidden Markov Directed Acyclic Graph (HMDAG) probabilistic model representing the collections of state spaces of time-stamped detection locations *𝒮*= U_*t*=1,…,*T*_ *𝒮*_*t*_ shown in the round nodes with conditional probability arrows depicting neighborhood structures and the observable outputs. The time-ordering constraint implies forward arrows only connecting groups of states to future time-stamped detection states. Notice the directed arrows denoting the conditional neighborhood Markov structure of descendents are conditionally independent given their parents nodes.

**Figure 5:**
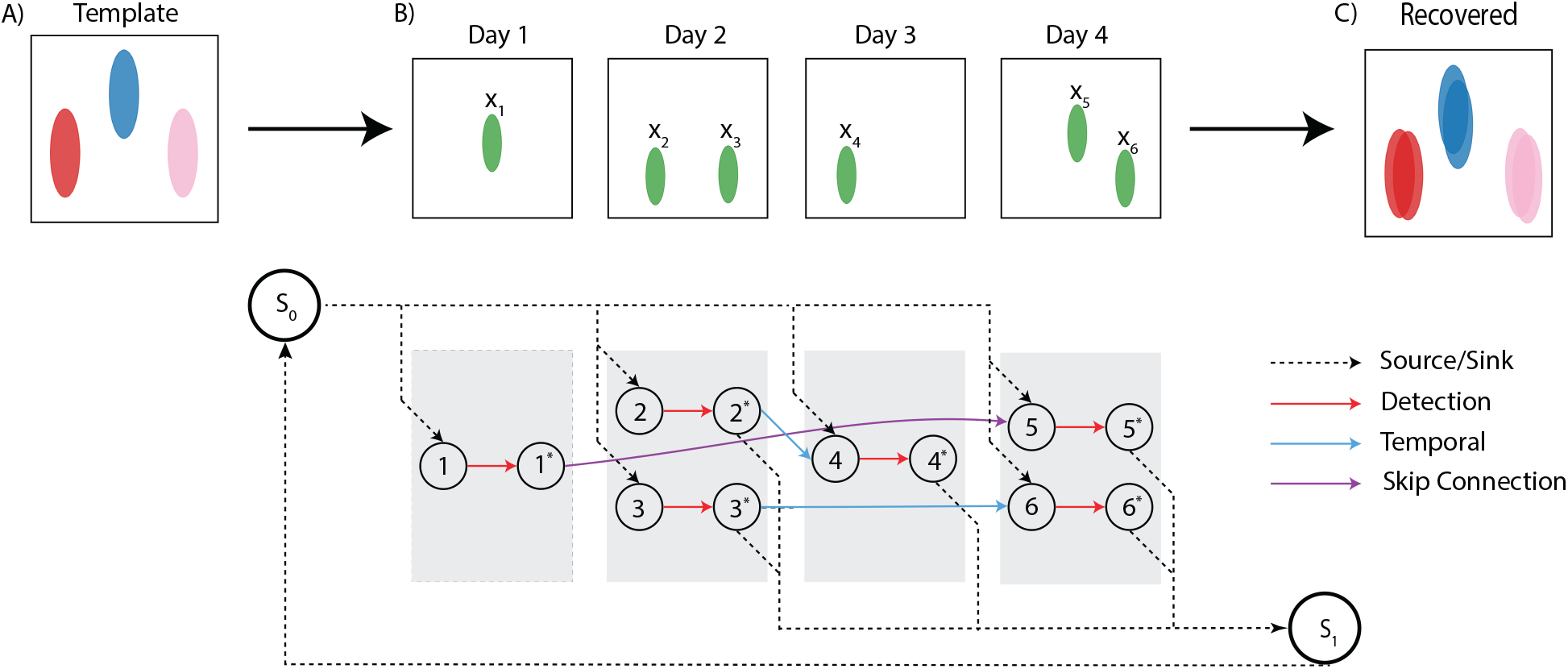
Example tracking problem with three fluorescence peaks. Left: the 3 true synapses in different colors being imaged in four sessions. Middle: Sequence of four imaging sessions with 1 detection on Day 1, 2 on Day 2, 1 on Day 3, and 2 on Day 4. Note that the detection at *x*_1_ disappears at Day 1 reappears at Day 4 at the location *x*_5_. Right: Cartoon reconstruction of the aligned tracks.

**Figure 6:**
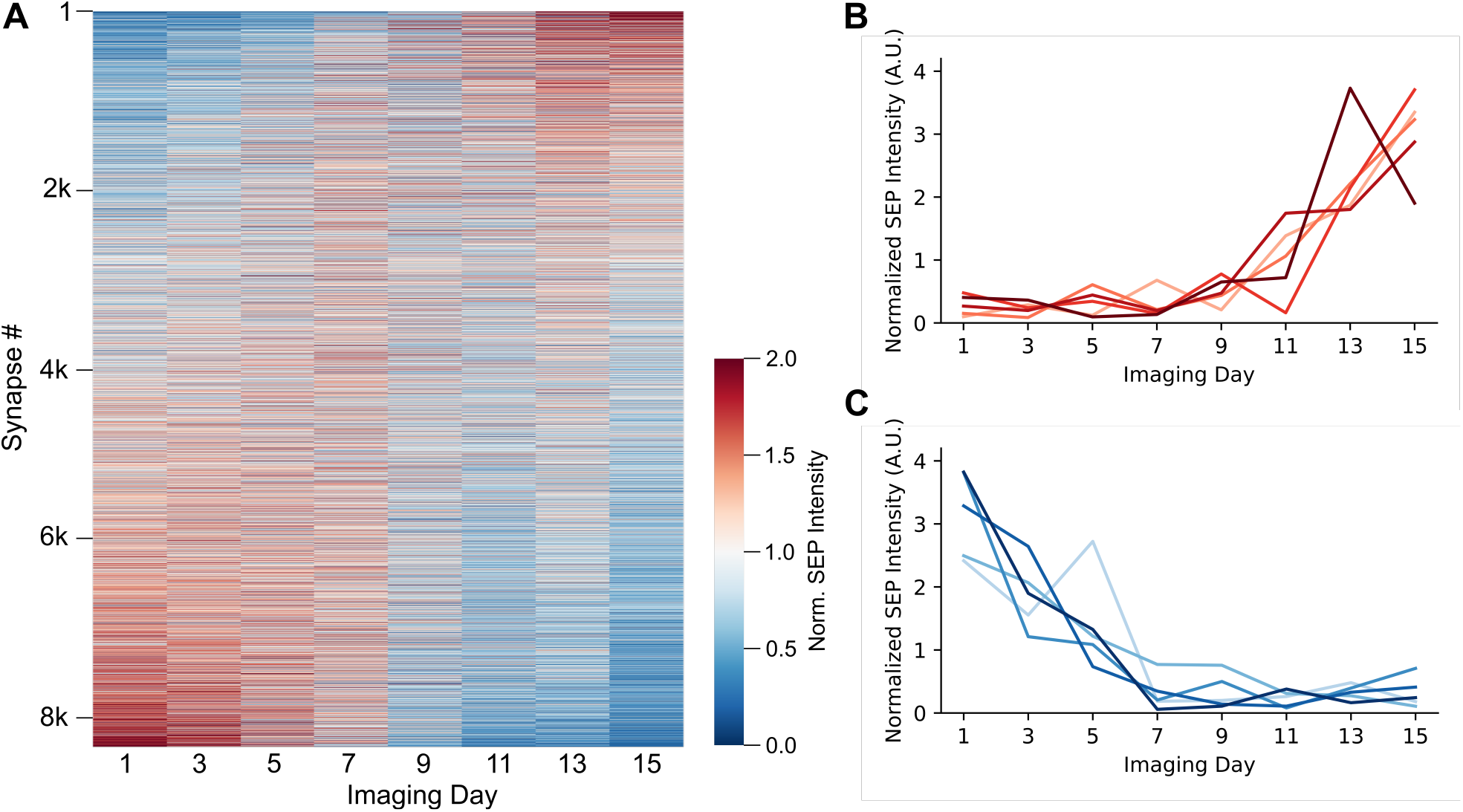
Synapse dynamics in vivo. (A) Normalized SEP-GluA2 intensity in a mouse cortex imaged over 2 weeks. Synapses were sorted by the slope of best fit line and each track’s SEP intensity was normalized to its average intensity. Heatmap shows synapses potentiating (top) and depotentiating (bottom). (B) Example traces from 5 potentiating synapses. (C) Example traces from 5 depotentiating synapses.

The CostScaling algorithm used by SynTrack has a polynomial computational complexity of 𝒪 (*n*^2^*m* log(*nC*)) [23] for a graph with *n* nodes and *m* edges and *C* being the largest edge weight. For efficiency, we used a C++ implementation of the Cost Scaling algorithm provided by the LEMON library. This resulted in a highly efficient tracker: running SynTrack on this dataset took roughly 10 minutes.

## 4 Discussion

We present SynTrack, an in vivo Synaptic Plasticity Tracking algorithm. Our results showcase the novel ability to accurately track changes in individual synaptic strength across dense networks of tens of thousands of synapses across two weeks in living mice. Together, these results provide, to our knowledge, one of the largest longitudinal measurements of synaptic plasticity at single-synapse resolution in vivo, revealing the dynamic processes of synaptic plasticity across large synaptic volumes over several weeks. Beyond this proof-of-concept demonstration, SynTrack can be used to track the synaptic foundations of any behavior by imaging any brain region of interest, provided sufficient optical access using in vivo two-photon microscopy. More broadly, the ability to accurately track longitudinal changes in fluorescent puncta intensity could be extended to other similar questions in neuroscience and biology. For instance, SynTrack could be used to follow changes in dendritic spine morphology and number to report structural plasticity associated with learning. Alternatively, these tools could be useful in tracking axonal boutons across either learning or development, or other fluorescently labeled proteins of interest, illuminating how complex biological systems form and remodel over time. As SynTrack’s accuracy is sensitive to upstream segmentation errors, future work should explore frameworks that jointly optimize detection, nonlinear registration, and tracking directly over raw image volumes, which could reduce error propagation across pipeline stages.

## Notes

* This work was supported by the National Institute of Neurological Disorders and Stroke under Grant R01NS134842

### Competing Interest Statement

The authors have declared no competing interest.

### Summary of Updates

We have updated and improved the algorithmic presentation and added additional comparisons.

